# Multivariate associative patterns between the gut microbiota and large-scale brain network connectivity

**DOI:** 10.1101/2020.08.25.266122

**Authors:** N. Kohn, J. Szopinska-Tokov, A. Llera, C. Beckmann, A. Arias Vasquez, E. Aarts

**Author notes:** Equal contribution (N. Kohn, J. Szopinska-Tokov, A. Arias Vasquez, E. Aarts). Corresponding author: Nils Kohn, Kapittelweg 29, 6525 EN Nijmegen, The Netherlands.

## Abstract

Research on the gut-brain axis has accelerated substantially over the course of the last years. Many reviews have outlined the important implications of understanding the relation of the gut microbiota with human brain function and behavior. One substantial drawback in integrating gut microbiome and brain data is the lack of integrative multivariate approaches that enable capturing variance in both modalities simultaneously. To address this issue, we applied a linked independent component analysis (LICA) to microbiota and brain connectivity data.

We analyzed data from 58 healthy females (mean age = 21.5 years). Magnetic Resonance Imaging data were acquired using resting state functional imaging data. The assessment of gut microbial composition from feces was based on sequencing of the V4 16S rRNA gene region. We used the LICA model to simultaneously factorize the subjects’ large-scale brain networks and microbiome relative abundance data into 10 independent components of spatial and abundance variation.

LICA decomposition resulted in four components with non-marginal contribution of the microbiota data. The default mode network featured strongly in three components, whereas the two-lateralized fronto-parietal attention networks contributed to one component. The executive-control (with the default mode) network was associated to another component. We found the abundance of *Prevotella* genus was associated to the strength of expression of all networks, whereas *Bifidobacterium* was associated with the default mode and frontoparietal-attention networks.

We provide the first exploratory evidence for multivariate associative patterns between the gut microbiota and brain network connectivity in healthy humans, taking into account the complexity of both systems.

## Introduction

The gut-brain axis (GBA) is the bidirectional biochemical signaling that takes place between the gastrointestinal tract (GI tract) and the central nervous system (CNS)(1). The microbiota-GBA is used to describe the complex effects of the commensal gut bacteria (the microbiota) in the interplay between the gut and the brain. Recently, many studies have outlined the important implications of understanding the relation of the gut microbiota with human brain function and behavior. Several intermediary pathways have been proposed: specifically, bi-directional interactions between microbiota and the brain are plausible via modulation of vagal nerve activity, via neuromodulators or their precursors such as serotonin or tryptophan, via the Hypothalamic-Pituitary-Adrenal System (HPA-axis) and via interactions with the immune system (1–4).

In recent years, researchers aimed at elucidating these interactions, highlighting putative pathways, hormonal or immunological agents, and targeting the activity and interaction of certain bacterial strains (3). However, these studies have not taken into account the complexity and, especially, the full multivariate nature of both the brain and the gut microbiome.

One of these complex traits of the brain is the intrinsic connectivity between different brain regions. So far, studies assessing the relation between gut microbiome composition and intrinsic brain connectivity – with resting state fMRI – are rare, limited in rigor, and inconclusive (5). A recent study tested the effects of four weeks multi-strain probiotics supplementation (6). The authors report mild probiotics-induced changes in resting state connectivity of some of the ten networks tested. The strongest modulation was found in differences between the placebo (n=15) and probiotics (n=15) group, with the latter showing a relatively stronger increase in connectivity of the salience network to superior frontal brain regions. In another placebo-controlled trial of probiotics (n=20 per group), *Bifidobacterium longum* influenced resting neural oscillations measured with magnetoencephalography (MEG), which correlated with enhanced vitality and reduced mental fatigue during a social stress induction task. Modulations of theta and alpha band oscillations by probiotics were localized in the frontal and cingulate cortex and supramarginal gyrus (7). However, these results (in relatively small samples) have not been related to probiotics-induced effects on gut microbiota composition.

A few studies did assess the relation between gut microbiome composition and intrinsic brain connectivity. One resting state fMRI study (n=30), which included a subgroup of smokers, focused on the association of gut microbiota composition with insula connectivity and found its connection to several brain regions, such as occipital and lingual gyrus, frontal pole and cerebellar regions, to be associated with microbiota diversity and structure (8). Other exploratory region-of-interest (ROI) analyses did not reveal significant associations. Another resting state fMRI study (n=28 vs 19) demonstrated that in end-stage renal disease, the integrity of the default mode network (DMN) was decreased along with alterations in the gut-microbiota composition (9). Taken together, these results are difficult to integrate and comprehend, as studies focus on one aspect of the modalities, such as connectivity from one particular ROI, or the gut-brain axis in targeted patient groups, or with different types of interventions. Most importantly, all previous studies have performed bivariate associations between one gut-microbiome composition measure and one brain connectivity measure (i.e. within one network or between two brain networks).

Indeed, in research, one approach to understand such complex systems is to try to elucidate the function of all its components sequentially and then to integrate interactions between a limited number of components. The opposite approach of investigation is to aim at integration at a macroscopic level. In this approach as many components as possible are sampled and patterns are investigated by dimensionality reduction. This has been attempted for the gut-brain axis very often narratively, in multiple reviews. Yet, no empirical attempt has been made so far to try to integrate the functions of the brain and the gut microbiome at a macroscopic level to determine associations between variance in macroscopic components of the two systems. In a similar approach, using data from the Human Connectome, researchers linked several lifestyle, demographic and psychometric measures in a positive-negative mode of brain connectivity (10).

Here, we aim to assess the relation between these two complex, multivariate modalities, focusing on canonically established brain networks in resting state that represent major modes of brain functioning in an unperturbed fashion (11). We asked if the inter-individual variability in abundance of gut microbiome genera was linked to variability in brain functional connectivity in canonical brain networks, when taking into account the full complexity of both.

One substantial methodological challenge is the multivariate and simultaneous integration of gut microbiome and brain data that enable capturing variance in both modalities simultaneously. To address this issue, we applied a linked independent component analysis ((12, 13), LICA) to microbiota and brain connectivity data (Figure 1). LICA enables data reduction in several modalities simultaneously and thereby is able to demonstrate joint inter-individual variation patterns in different modalities. We chose to investigate four very well characterized and often replicated brain networks (11). We used this selection in previous work to investigate the impact of fasting on functional connectivity in rest (14). We limited our study to a set of four networks of interest (the lateralized fronto-parietal (left/right) attention networks, FPN; the executive control network, ECN; and the default mode network, DMN) due to their importance in the neuroimaging field, their comparatively clear and cognitive functional profile and their importance in mental disease or previous microbiome research (10, 11, 14–16).

**Figure 1.**
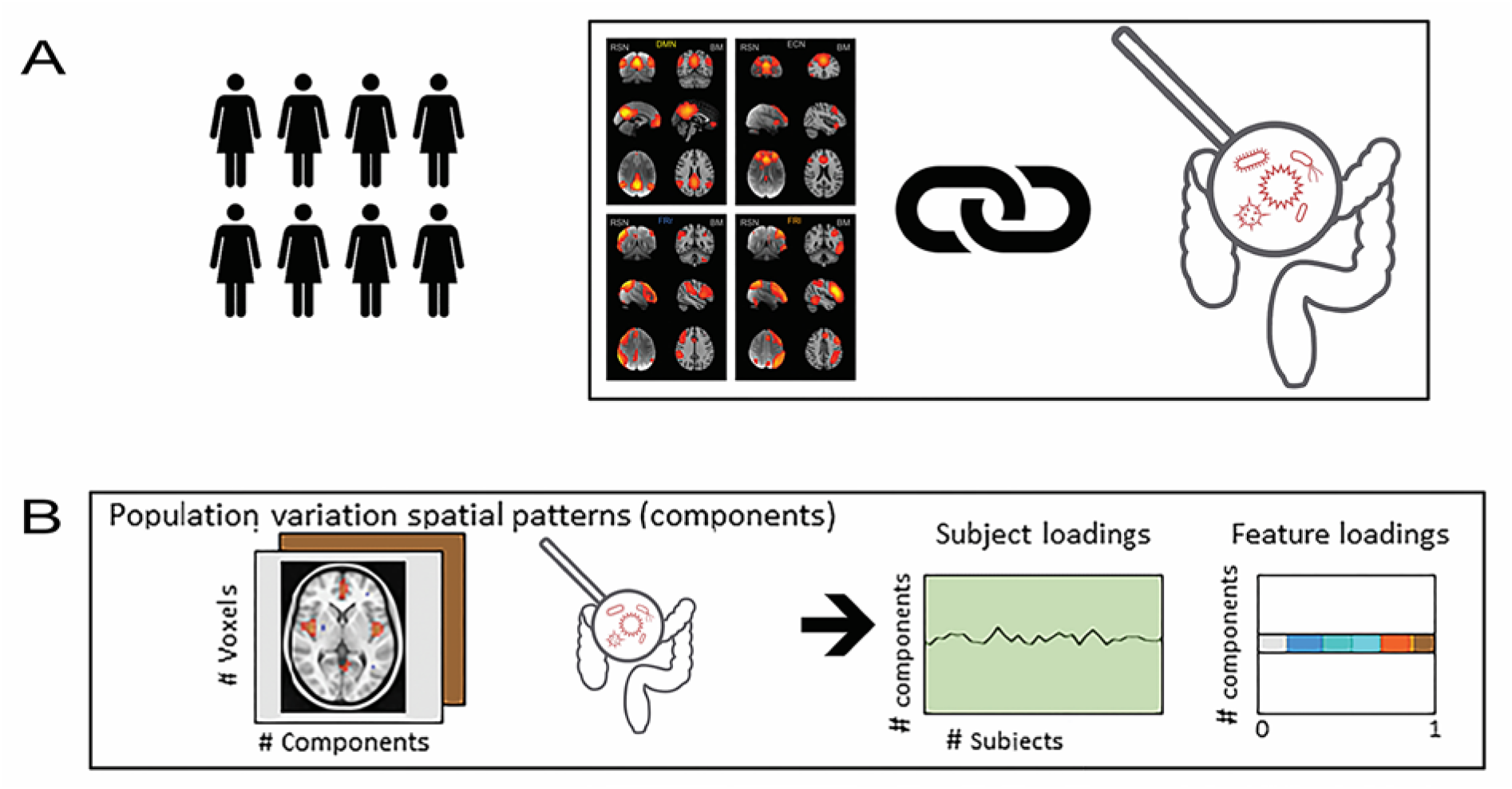
We linked functional brain connectivity in four well-established brain networks with relative abundance of human gut bacteria (microbiota). Panel A describes the Linked ICA that decomposed, simultaneously, the variability in functional connectivity of the four networks and the relative abundance of bacterial taxa (genera). This resulted in 10 components for which we have individual subject loadings as well as the loadings of each input feature depicted in panel B. The loadings represent voxel-wise association to the component in functional connectivity per network and genera-wise association to the component in the gut-microbiome.

## Results

For 58 subjects, the spatial template maps of right and left frontoparietal-attention networks (FPN), executive control (ECN), default mode network (DMN) (11) were projected onto the subjects resting state fMRI time-courses to create network maps per subject. Gut microbiome composition was based on sequencing of the V4 region of the 16S rRNA gene on the Illumina HiSeq platform. We used the LICA model to simultaneous factorize the subjects’ brain networks and gut microbiome relative abundance into ten independent components (12, 13).

### Joint decomposition of brain networks and microbiome relative abundance

From the 10 components, six showed a non-marginal (proportion > 0.2) contribution on both the gut microbiota relative abundance and the brain connectivity patterns (Component 0, 1, 3, 4, 6 and 7ö Figure 2). From these six components, the first extracted component (component 0) was explained by a single subject, therefore, this component was disregarded for further analyses. Additionally, sanity checks on brain connectivity showed, for component 4, equal values for all voxels in the brain data. This renders interpretation of this component hardly possible and could potentially relate to residual noise being picked up and explained. This component was therefore also discarded.

**Figure 2.**
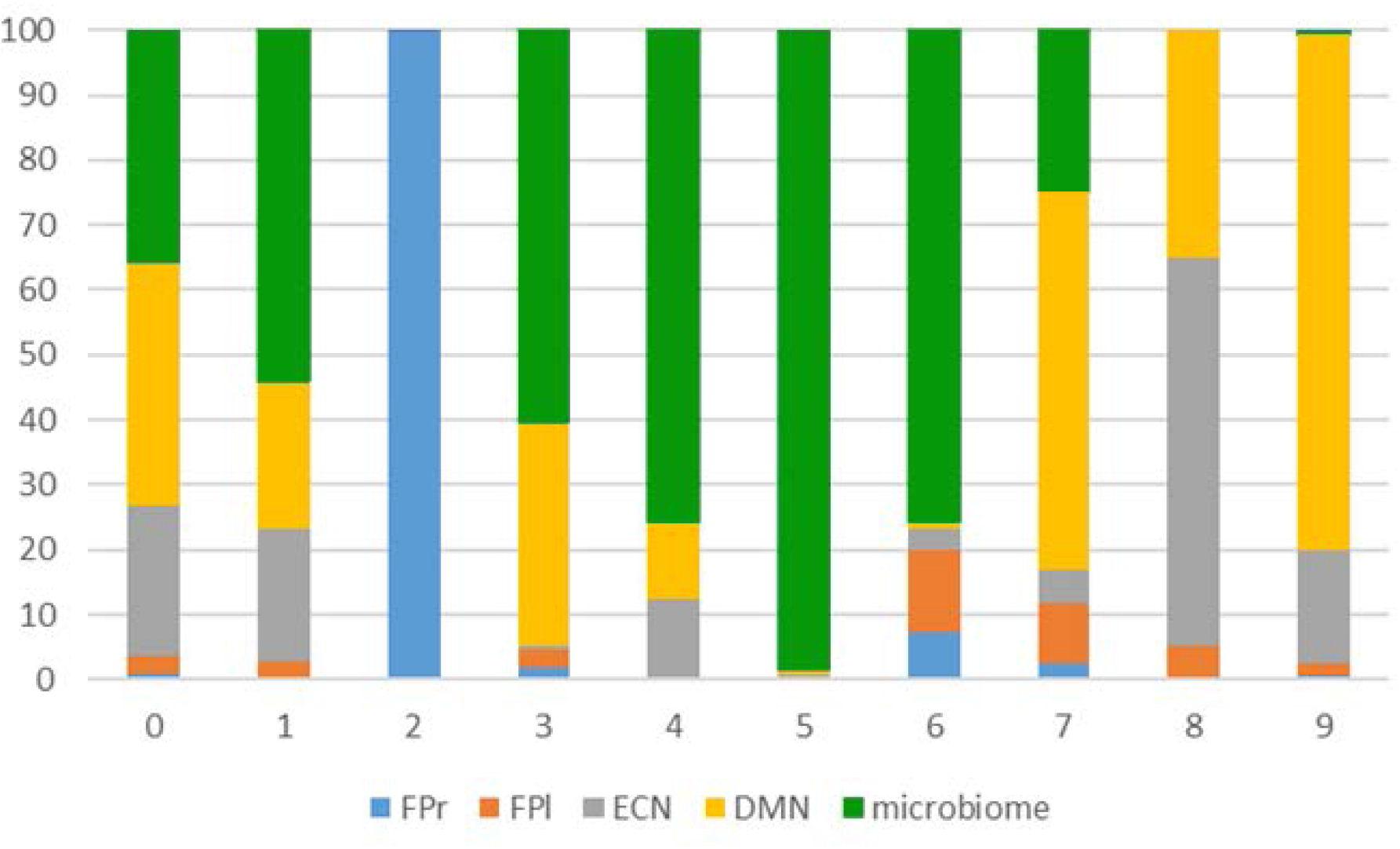
Decomposition of brain connectivity and microbiome. The plot shows the percentage of contribution per input modality. FPr and FPl are right and left fronto-parietal networks, DMN is default mode and ECN is executive control network.

We consequently investigated the association between brain connectivity and gut microbiota relative abundance in the four remaining components. We characterized each component by the contribution of the different modalities (proportion > 0.2 for brain or microbiome). For each component, we plotted the brain-network and their voxel-wise loading and listed the bacterial genera that were non-marginally associated with the component. For brain connectivity data, the z-maps from the Linked ICA were thresholded at a z > 3 for display purposes (see Neurovault: https://neurovault.org/collections/TRVFBPAB/ for z maps of the brain data four components). The z-scores reflect how strongly a voxel covaries in connectivity with the respective input network. For microbiome data, high loadings reflect a robust covariation in relative abundance of a particular genus in that particular component. Similarly, we thresholded the microbiome loading at z>2.3 as they were more sparse compared to brain loadings (See Figure 3 for a visualization of these results and the DondersSharingCollection for the unthreshholded decomposition data).

**Figure 3.**
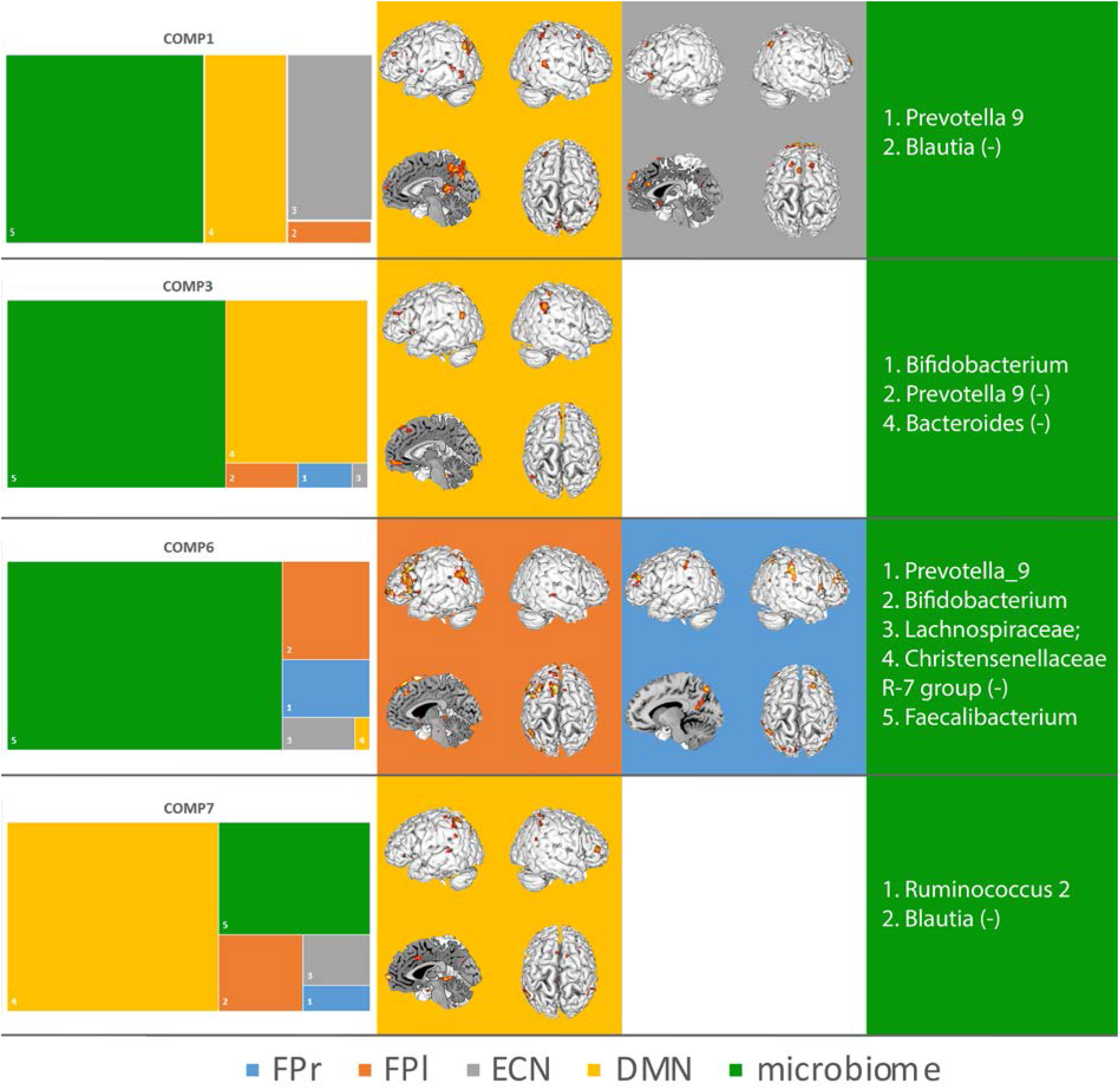
Summary results of the contribution of each modality are shown in the first column (left). Second and third columns display the spatial project of the brain modalities, e. g. which voxels covary most strongly with covariation in other modalities (brain networks and microbiota abundance). Fourth column (right) displays the genera that show a covariance in abundance that is linked to covariance in the brain networks. The colors align with the modalities of the LICA (the four brain networks and the gut microbiota). For display purposes genera loading were cut at z >2.3 (for more details see the method section). See supplementary material for full list of genera loadings on the four components.

The microbiota accounted for the majority of the variability that could be explained by component 1, 3 and 6. Component 1 also has strong contributions from variability in the DMN and the ECN and the microbiota. Looking at the variability in each modality more deeply, the LICA enabled us to say which voxels show variation in functional connectivity to the respective network between subjects. For the microbiota, LICA gives an indication of between-subject variation in relative abundance of the gut microbiota genera for that component. For instance, ECN in component 1 varied between subjects in core hubs of the ECN, such as the dorsal paracingulate cortex, middle frontal and superior frontal gyri. The ECN largely consists of middle frontal and superior frontal gyri, paracingulate cortex and dorsal posterior parietal cortex (11, 17). This covariance in core hubs of the ECN can be interpreted as this component explaining functional connectivity strength, or the strength of the expression of the ECN in the subjects, and this strength of expression being related to variation in relative abundance of gut microbiota. The ECN has been demonstrated to overlap spatially with brain activity observed in cognitive control tasks, emotion tasks and response inhibition (11). Similarly, for the DMN, component 1 picked up on variability in the posterior core hub of the DMN (the posterior cingulate and retrosplenial cortex; 13, 16). The DMN is the large-scale brain network that was identified first and it is probably the most often studied of all so-called resting-state networks. DMN modulations have been implicated in a broad range of disorders (19–22). The most prominent feature of the DMN is its task-negative nature; the areas of the DMN deactivate when an individual is engaged in most tasks (23–25). It has been associated to a broad range of cognitive processes such as self-referenced thought and self-monitoring (23), passive, broad attention (20, 26), auto-biographical memory retrieval (27, 28), imparting meaning in the current sensory input depending on prior experiences (29), mind-wandering and future thinking (29, 30) as well as homeostatic functions (23, 24, 31, 32). *Prevotella_9* was more abundant and *Blautia* was less abundant with increasing between-subject functional connectivity of these hubs of the two networks. Component 2 had a contribution of over 50% from variability in microbiota abundance and the DMN. *Bifidobacterium* was more abundant and *Prevotella_9* and *Bacteroides* were less abundant with increasing functional connectivity in anterior core hubs of the DMN. Component 6 had the strongest contribution of all components from microbiota of around 75%, yet also explained variability in the two-lateralized fronto-parietal attention networks. As the name of these networks suggest, they encompass fronto-parietal brain regions, which are commonly and reliably associated to brain activity in attention tasks (11) and are modulated with varying degree of attention demand (33). The topology of the loading of these networks on this component overlaps with their common, canonical spatial profile in lateral frontal and parietal brain areas. Thus, again this component is associated to the between-subjects variation of the strength of expression of the lateralized attention networks. *Prevotella_9, Bifidobacterium*, genera belonging to *Lachnospiracaceae* family, and *Faecalibacterium* were more abundant and *Christensenellacea_R-7_group* was less abundant with stronger expression of the attention networks. Component 7 was associated to variability in the DMN again and to roughly 25% of the microbiota. The spatial pattern of between-subject variation could be interpreted as elevated connectivity of the DMN to parts of the so-called salience network (such as dorsal anterior cingulate cortex (dorsal ACC) and ventrolateral prefrontal cortex (VLPFC)), which has in previous literature been associated with effects of elevated stress on DMN resting state connectivity (15, 16). *Ruminocuccus_2* was more abundant and *Blautia* was less abundant in individuals that showed this elevated connectivity pattern of DMN to dorsal ACC and VPLFC.

## Discussion

In this study, we provide the first evidence for multivariate associative patterns between the gut microbiota and brain network connectivity in healthy humans. We used a novel multivariate modality integration technique to explain inter-individual differences in brain connectivity in four canonical networks and the gut microbiota. We see our exploratory results as a map that could show high potential to guide future research on the relation of gut-brain interactions in a hypothesis-generating manner. We have linked ECN connectivity to an abundance of *Prevotella_9 and Blautia;* DMN connectivity to *Prevotella_9, Blautia, Ruminococcus_2, Bifidobacterium*, and *Bacteroides;* frontoparietal attention network connectivity to *Prevotella_9, Bifidobacterium, Faecalibacterium, Christensenellacea_R-7_group*, and certain genera belonging to *Lachnospiracaceae*. DMN connectivity that has been linked to stress is associated with *Ruminocuccus_2 and Blautia*. The spatial associations in the components to core hubs of the respective networks can be seen as a conceptual validation of our approach (11, 16, 34). Furthermore, we observe the between-subject variation in functional connectivity in core hubs of the respective networks in three of the four components as a link between an individual’s connectivity strength and the relative abundance of certain microbiota. These findings can be taken as an indication that certain microbial genera are associated with the normal expression of all four canonical resting state networks and their natural variation between healthy subjects.

On the side of the bacterial genera that were associated with brain network connectivity, we found that inter-individual variation in abundance of the *Bifidobacterium* genus was prominently contributing to two of our four identified components. Variation in abundance of *Bifidobacteria* were associated with increased connectivity of the medial prefrontal cortex of the DMN and parietal regions and in another component with modulated connectivity of the core hubs of the fronto-parietal attention network. The *Bifidobacterium* genus is probably one of the most noticeable targets in current gut-brain axis research (2, 35, 36). This strong focus is potentially related to a landmark study, which showed that germ-free mice have altered HPA-axis function, and this altered HPA activity was reversed by colonization with a *Bifidobaterium* (1). *Bifidobacteria* are one of the most important and abundant genera during development and have been associated with decreased levels of inflammation in human development (37). *Bifidobacterium longum*, a strain commonly used in probiotic products, influenced resting neural activity that correlated with enhanced vitality and reduced mental fatigue during a social stress induction task (7). The medial prefrontal cortex and the DMN have been related to autobiographic and episodic memory or prior knowledge structures(38), which fits to findings of the link between increased *Bifidobacteria* after interventions and elevated verbal episodic memory(6). Furthermore, Bagga and colleagues also found altered functional connectivity of the DMN after probiotic use (including *B. longum*) (6). Component 3 might therefor partially reflect episodic memory-related modulations in *Bifidobacteria* and DMN connectivity. A probiotics trial with *Bidfidobacterium longum* using electrophysiological resting state brain recordings found evidence for an association of increased frontal midline mobility and improved memory after probiotics consumption compared to placebo (39). The authors related the brain recordings to attention-related brain activity. Moreover, the fact that out of >50 genera that featured in our analysis, *Bifidobacteria* featured in two of the four components both underscores their putative influence in the gut-brain interaction and the validity of our integrative approach. In summary, we found evidence for a relation of *Bifidobacteria* abundance to attention- and potentially memory-related brain network activity at rest.

For component 7, the spatial patterns of association were similar to results showing an alerted state of the DMN after social stress induction (15). In data using a similar paradigm, *Bifidobacterium longum* modulated activity in similar regions that were influenced by social stress and also in the hippocampus, a region that is part of the DMN (7). This pattern particularly varied with abundance in *Blautia* and *Ruminococcus 2*. Although *Bifidobacterium* did not covary with this component, the *Bifidobacterium* intake might have indirectly affected DMN connectivity in stress, potentially via modulation of the abundance of *Ruminococcus 2* and *Blautia*. Indeed, the pre-existing levels of *Blautia* and *Ruminococcae* correlated with the metabolic outcomes of a *Bifidobacterium-targeting* prebiotic intervention in obese patients(40). Furthermore, *Blautia* has been found to be the only genus to be enriched in depression-model rats(41) and both *Blautia* and *Ruminococcae* correlated with stress-related depression-like behaviour in mice(42). In summary, variations in *Blautia* and *Ruminococcus 2* abundance might relate to stress-induced modulation of DMN connectivity.

The association of Components 1, 3 and 6 with *Prevotella_9* is interesting as this genus has been previously involved in psychiatric disorders, cognition and brain connectivity changes. For example, in autism spectrum disorder (ASD), which is characterized by atypical brain network organization (including DMN and ECN, as in component 1) (43), a higher relative abundance of *Prevotella* (and *Bifidobacterium*) has been linked with a beneficial effect of Microbiota Transfer Therapy (44). Accordingly, lower relative abundance of *Prevotella* has been associated with psychiatric disorders like ADHD in children (45), Parkinson’s disease (46), and ASD (47). Furthermore, the gut-brain axis may play a role in the disturbed executive functioning in ASD (for a review, see (48)). Our finding of a positive correlation of *Prevotella* with DMN and ECN functioning and also fronto-parietal attention network modulations support these results. Previous work showed a link between gut microbiota and resting-state functional connectivity, as assessed here (9, 49). Interestingly, in one study assessing bivariate relationships, *Prevotella* and *Bacteroides* were associated with insular connectivity (8). The insula has not only been discussed as part of the salience network, but also as an important component of the general task positive network (50, 51). In our case, both of the *Prevotella* and *Bacteroides* genera were negatively associated with DMN in component 3. As the DMN is thought to be anti-correlated with the task positive network, our finding is in line with previous results (50). *Prevotella* seem to be associated to healthy modulation in brain connectivity related to attention, cognitive control, episodic memory and a range of other psychological functions.

Our design and approach have limitations in the interpretation of the results. First, these findings are necessarily limited to more common genera. We capped our analysis at genera that are at least detectable in 30% of our subjects. Genera with lower occurrence rates in individuals might have unequally strong leverage on the LICA. As a consequence of this methodological choice, we cannot exclude an overestimation of the loadings for the more common taxa (given the sample size) and we cannot assess the rare genera and their association with brain network connectivity.

Second, the selection of brain networks was motivated by their role in cognition specifically to high-level cognitive constructs such as attention and cognitive control and their relevance in the literature. While we perceive this selection as well-motivated and we have demonstrated their sensitivity (14), it is a subjective pre-selection. We might not cover other cognitive processes and associated brain networks equally well. Nevertheless, due to the limited power in our sample and for the advantage of choosing networks that are more readily interpreted, we chose to limit our selection of brain networks to these four.

Third, we investigated a very homogenous, healthy and young group of only female participants. Although this naturally limits the generalizability of the results, we believe that our data still serves an orientating purpose and is therefore valuable. In replication attempts, this homogeneity and special characteristic of our sample should be considered. We would like to reiterate that we see a strong need to replicate the current results in larger and more diverse samples.

In summary, we provided the first evidence for multivariate associative patterns between large-scale brain network functional connectivity of four very well-established brain networks and the relative abundance of gut microbiota in a sample of healthy female individuals. This link provides a map for future research, involving the full complexity of both measures into account. For example, interventions targeting improvement in attention (for example in neurodevelopmental disorders) could investigate the influence on the bacterial genera associated to the attention networks. Moreover, it can provide a roadmap to investigate how the effect of probiotic intervention trials with compounds that benefit certain genera could explicitly test modulations in cognitive functions associated to the brain networks we investigated or to brain connectivity itself. Furthermore, future research might investigate the mechanistic nature of our multivariate associative patterns and aim to assess the generalizability to other healthy samples as well as their potential disruption in the diseased brain.

## Material and Methods

### Sample

We analyzed pre-intervention data from a probiotics intervention study on 64 healthy female participants (mean age = 21.5 (0.45) years) (52). In total, 58 of the 64 participants were included in the analyses. Six participants were excluded from the final analyses, due to high depression scores (N=1), missing feces samples (N=2), and movement exceeding 4mm between acquisitions (n=3). For more detailed characteristic of the samples and exclusion criteria as well as the ethical declaration, please see the Material and Methods section of Papalini et al. (52). Briefly, participants with relevant medical history of e.g. psychiatric and/or gastrointestinal disorder were excluded. Also, use of antibiotics and diet like e.g. vegan diet were part of the exclusion criteria.

### fMRI data acquisition

Participants were screened for compatibility to magnetic resonance imaging (MRI). MRI data were acquired using a 3T MAGNETOM Prisma system, equipped with a 32-channel head coil. After three short task-related fMRI scans (see Papalini et al.), nine minutes of resting state fMRI was acquired. 3D echo planar imaging (EPI) scans using a T2*weighted gradient echo multi-echo sequence (Poser, Versluis et al. 2006) were acquired (voxel size 3.5 x 3.5 x 3 mm isotropic, TR = 2070 ms, TE = 9 ms; 19.25 ms; 29.5 ms; 39.75 ms, FoV = 224mm). The slab positioning and rotation (average angle of 14 degrees to AC axis) optimally covered both prefrontal and deep brain regions. Subjects were instructed to lie still with their eyes open and refrain from directed thought. A whole-brain high-resolution T1-weighted anatomical scan was acquired using a MPRAGE sequence (voxel size 1.0 x 1.0 x 1.0 isotropic, TR = 2300 ms, TE = 3.03 ms, 192 slices).

MRI data preprocessing: FSL (FMRIB, University of Oxford, UK; www.fmrib.ox.ac.uk/fsl; (Jenkinson et al., 2012) was used for pre-processing, data-denoising, and generation of subject-specific network maps. Pre-processing steps included three-dimensional movement correction, and spatial smoothing using a 5 mm full-width at half maximum (FWHM) Gaussian kernel to reduce inter-subject variability and a high-pass filter (> 0.007 Hz) was applied. All pre-processing steps, except temporal filtering, were conducted before AROMA data denoising (53, 54). Briefly, ICA-AROMA is designed to identify motion-related artifacts by matching single subject ICA components to four robust and standardized features. The data is denoised by linear regression of ICA components identified as noise by AROMA and subsequently the high pass filter was applied. Prior to all group analyses, data were normalized to MNI space and re-sampled to 2 mm^3^ resolution using FMRIB’s Nonlinear Image Registration Tool (FNIRT).

### Generation of subject-specific functional connectivity maps

Dual (spatial and temporal) regression was used to generate subject-specific spatial maps of well-studied, canonical large-scale brain networks(11) from the individuals’ data. The z-maps of these networks were temporally concatenated in one 4D file and used as input for the dual regression. These maps were used in a linear model fit against the individual fMRI data, resulting in the subject-specific temporal dynamics. Subsequently, these time-course matrices are employed in a linear model fit against the subject’s fMRI data set to estimate subject-specific spatial maps. From these subject-wise expressions of the 10 networks, we selected four networks of interest (the left and right lateralized fronto-parietal attention networks, FPN; the executive control network, ECN; and the default mode network, DMN), due to their importance in the neuroimaging field, their comparatively clear functional profile and their importance in mental disease or previous microbiome research (10, 11, 14–16). The different spatial maps for all participants are combined into a single 4D file per target network. In this way, we generated four files for the four respective networks of interest that contain one spatial z-map per subject that indicates for each voxel the connectivity strength of the respective network in that individual. These four network files were used as inputs to the Linked-ICA.

### Gut microbiome analysis

Fecal samples were collected by using OMNIgene•GUT kit (DNAGenotek, Ottawa, CA) (55). Collected fecal samples were transported to the laboratory and aliquoted into 1.5 mL Eppendorf tubes and stored at −80 °C for microbiome analysis. DNA was isolated from the fecal pellets using the Maxwell^®^ 16 Instrument (Promega, Leiden, The Netherlands) as described previously (56). Briefly, in the 2-step PCR protocol the 16S rRNA gene V4 variable region was targeted by using 515F (GTGYCAGCMGCCGCGGTAA) and 806R (GGACTACNVGGGTWTCTAAT) primers, and unique barcodes were used to characterize a mixture of bacteria. Sequencing was performed on the Illumina HiSeq PE300 platform by GATC Biotech AG (Konstanz, Germany). The sequences were processed using NG-Tax (57) analysis pipeline as described previously (58). This resulted in an Operational Taxonomical Unit (OTU) table containing 844 OTUs. We applied a prevalence-filtering at the genus level, selecting genera present in at least 30% of the samples. After this step, the OTU-table containing 644 OTUs was used for the downstream analyses. The gut microbiome composition tables at the phylum and genus taxonomic levels were provided by the ‘phyloseq’ package available in R (59).

### Linked analyses

We used the Linked-ICA model (13) to simultaneous factorize the functional network maps (of ECN, FPNs and DMN) and the microbiome data of 58 subjects into independent sources (or components) of variation. In the brain networks, spatial variation was explained; while in the microbiome data, variation in relative abundance of bacterial genera was explained. In brief, Linked-ICA is an extension of Bayesian ICA (60) to multiple input sets, where all individual ICA factorizations are linked through a shared common mixing matrix that reflect the subject-wise contribution to each component (Figure 1).

This operation is represented in Figure 1. Factorization provides a set of spatial maps (one per feature modality and component), a vector of feature loadings that reflects the degree to which the component ‘represents’ the different modalities, and a vector that reflects the contribution of the individual subject to a given component. All mathematical derivations involved in the Linked-ICA factorization can be found in the original paper describing the original algorithm (13). Further details and code implementing each feature extraction procedure as well as the Linked-ICA factorization are publicly available at (61). Given the sample size, we forced a 10 components solution. We disregarded components estimated with marginal (proportion < 0.2) contribution of the microbiome or brain networks, respectively.

Data will be shared via the DondersSharingCollection (link to follow).

## Acknowledgements

The study was supported by the Dutch Ministry of Economic Affairs under the TKI Life Science and Health, project LSHM15034. CB and AL have received funding from the Innovative Medicines Initiative 2 Joint Undertaking under grant agreement No 777394 for the project AIMS-2-TRIALS. This Joint Undertaking receives support from the European Union’s Horizon 2020 research and innovation programme and EFPIA and AUTISM SPEAKS, Autistica, SFARI. This work was further supported by the European Union Horizon2020 programme CANDY (Grant Agreement No. 847818). EA received funding from the European Research Council (ERC_StG2019_852189). JST and AAV have received support from the European Union’s Horizon 2020 research and innovation programme under grant agreement No 728018 (Eat2beNICE).

